# The Effect of a Virtual No-Step Zone on Balance Control in Walking

**DOI:** 10.1101/2020.02.28.970152

**Authors:** Tyler Fettrow, Stephen DiBianca, Fernando Vanderlinde dos Santos, Hendrik Reimann, John Jeka

## Abstract

Humans need to actively control their upright posture during walking to avoid loss of balance. We do not have a comprehensive theory for how humans regulate balance during walking, especially in complex environments. Balance must be maintained in a variety of contexts including crowded city side-walks, rocky nature trails, walks on the beach, or fast-paced sporting events. The nervous system must process many aspects of the environment to produce an appropriate motor output in order to maintain balance on two legs. We have previously identified three balance mechanisms that young healthy adults use to maintain balance while walking: 1) The ankle roll mechanism, a modulation of ankle inversion/eversion; 2) The foot placement mechanism, a shift of the swing foot placement; and 3) The push-off mechanism, a modulation of the ankle plantarflexion angle during double stance. We know that these mechanisms are interdependent and can be influenced by internal factors such as the phase of the gait cycle and walking cadence. Here we seek to determine whether there are changes in neural control of balance when walking in the presence of environmental constraints. Subjects walked on a self-paced treadmill while immersed in a virtual environment that provides three different colored pathways. Subjects were instructed not to step in the *No-Step Zone*, which appeared either on the right or left side of the subject. While walking, subjects received balance perturbations in the form of galvanic vestibular stimulation, providing the sensation of falling sideways, either toward the *No-Step* zone or toward the *Neutral* zone on the other side. The results indicate that the use of the balance mechanisms are subtly altered depending on whether the perceived fall is toward the *No-Step* or the *Neutral* zone. This experiment provides further evidence that the balance control system during walking is extremely flexible, recruiting multiple mechanisms at different times in the gait cycle to adapt to environmental constraints.

## 1. Introduction

Human balance control has been extensively studied in standing (Winter et al., 1998; Kiemel et al., 2008; Peterka, 2002), but a comprehensive theory for how it is accomplished during walking is still not developed. Compared to standing, the gait cycle adds a layer of complexity to bipedal systems during locomotion. Balance while walking is further complicated in a dynamic and variable environment in walking compared to standing. Balance must be maintained in a variety of contexts, such as crowded city side-walks, rocky nature trails, walks on the beach, or fast-paced sporting events. How do environmental constraints contribute to the required motor output in order to maintain bipedal balance? Our interest here is to investigate a paradigm that systematically alters the use of the balance mechanisms during walking in response to constraints in the environment.

To our knowledge it is not understood how balance is achieved in the presence of environmental constraints. The majority of literature surrounding environmental constraints focuses on stepping (Rietdyk and Rhea, 2011) or steering around (Patla et al., 1991) an obstacle. These paradigms are valid representations of environmental constraints, as slipping and tripping combined account for an estimated 46% of falls among older adults (Leavy et al., 2015). If environmental constraints are imposed, we must adapt, and decide if falling in a certain direction has increased consequences or costs. Obviously any fall is undesirable, but certain types of falls may be more harmful, such as falling off a curb or a flight of steps.

A real-world situation that may alter the method of balance, although extreme and probably rare, could be walking along a narrow mountain foot-path with a 100m sheer cliff on the left side. A sudden gust of wind or a trip shifts the center of mass to the left. What does the nervous system do? A step to the left may be costly as this could shift the base of support to an unstable portion of the trail and result in a fall. In this example, we argue the nervous system must choose a specific balance response that reflects the cost of falling in a particular direction.

We know that the healthy human nervous system, without any environmental constraints, typically uses three major balance mechanisms (Reimann et al., 2018): 1) The ankle roll mechanism, a modulation of ankle inversion/eversion that shifts the center of pressure under the stance foot towards the perceived fall, pulling the body in the opposite direction; 2) The foot placement mechanism, a shift of the swing foot placement towards the perceived fall that shifts the center of pressure towards the perceived fall after heel-strike; and 3) The push-off mechanism, a modulation of the ankle plantarflexion/dorsiflexion angle of the trailing leg, accelerating the CoM not only in the anterior-posterior direction, but also in the medial-lateral direction to correct the perceived fall. The nervous system combines these mechanisms in different ways to maintain balance, as the use of each mechanism varies on any given step (Fettrow et al., 2019b). Furthermore, the relative use of the mechanisms is altered based on internal constraints, such as the phase of the gait cycle (Reimann et al., 2019), or the stepping cadence (Fettrow et al., 2019a). This allows for a highly flexible system that can meet the goal of maintaining balance during locomotion through a variety of scenarios.

Here we test whether the recruitment of the mechanisms depends on the perceived availability of places to step in the environment. It is known that vision is used to assign constraints to the environment in which the person is navigating (Patla et al., 1991; Patla and Greig, 2006; Jansen et al., 2011) and for planning of future step location (Matthis and Fajen, 2014). We created a paradigm that attempts to recreate the cliff trail example, without the factor of fear. The literature has focused on the use of the foot placement mechanism in order to maintain balance (Townsend, 1985; Bruijn and Van Dieën, 2018). We know the foot placement is actively modulated in response to a perceived fall (Hof and Duysens, 2013; Rankin et al., 2014; Reimann et al., 2017), and during normal walking (Wang and Srinivasan, 2014). If the foot placement mechanism is constrained, other mechanisms must be used in order to respond to a perceived threat to balance. We attempted to achieve this by placing “No Step Zones” lateral to the subject’s walking path, and periodically providing sensory perturbations to balance, providing the sensation of falling to the side, either toward the *No-Step* zone or the *Neutral* zone. We hypothesize that the foot placement mechanism will be restricted when the balance perturbation produces a perceived fall towards the *No-Step* zone.

## 2. Methods

20 healthy young subjects (13 female, 23.6 ± 4.48 years, 1.68 ± 0.096 m, 70.14 ± 10.48 kg) volunteered for the study. Subjects provided informed verbal and written consent to participate. Subjects did not have a history of neurological disorders or surgical procedures involving the legs, spine or head. The experiment was approved by the University of Delaware Institutional Review Board (IRB ID: 1339047-4).

### 2.1. Experimental design

Figure 1 shows the experimental setup, with subjects walking on a split-belt, instrumented treadmill (Bertec, Columbus, OH, USA, Figure 1A) within a virtual environment displayed via an Oculus Rift (Facebook Inc., Menlo Park, CA, USA, Figure 1B). The treadmill was self-paced, using a nonlinear PD-controller in Labview (National instruments Inc., Austin, TX, USA) to keep the middle of the posterior superior iliac spine markers on the mid-line of the treadmill. The head position in the virtual world was linked to the midpoint between the two markers on top of the Oculus headset, in which optic flow (forward motion) was defined by the treadmill speed. Rotation in the virtual world was updated via the IMU embedded in the Oculus. We adjusted yaw drift of the IMU through use of real-time rigid body 6DOF tracking of the head via Qualisys (Gothenburg, Sweden) motion capture software, according to the equation

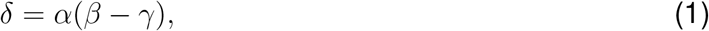

where *β* is the yaw angle of the head-mounted display rigid body from motion capture, *γ* the internal yaw angle of the Oculus IMU and *α* a gain factor. This equation corrected the internal yaw angle of the Oculus system to the motion capture value by a small factor each time step, effectively tethering the Oculus perspective to the motion capture to remove long-term drift from gyroscope integration. The gain parameter was set to *α* = 0.05.

**Figure 1:**
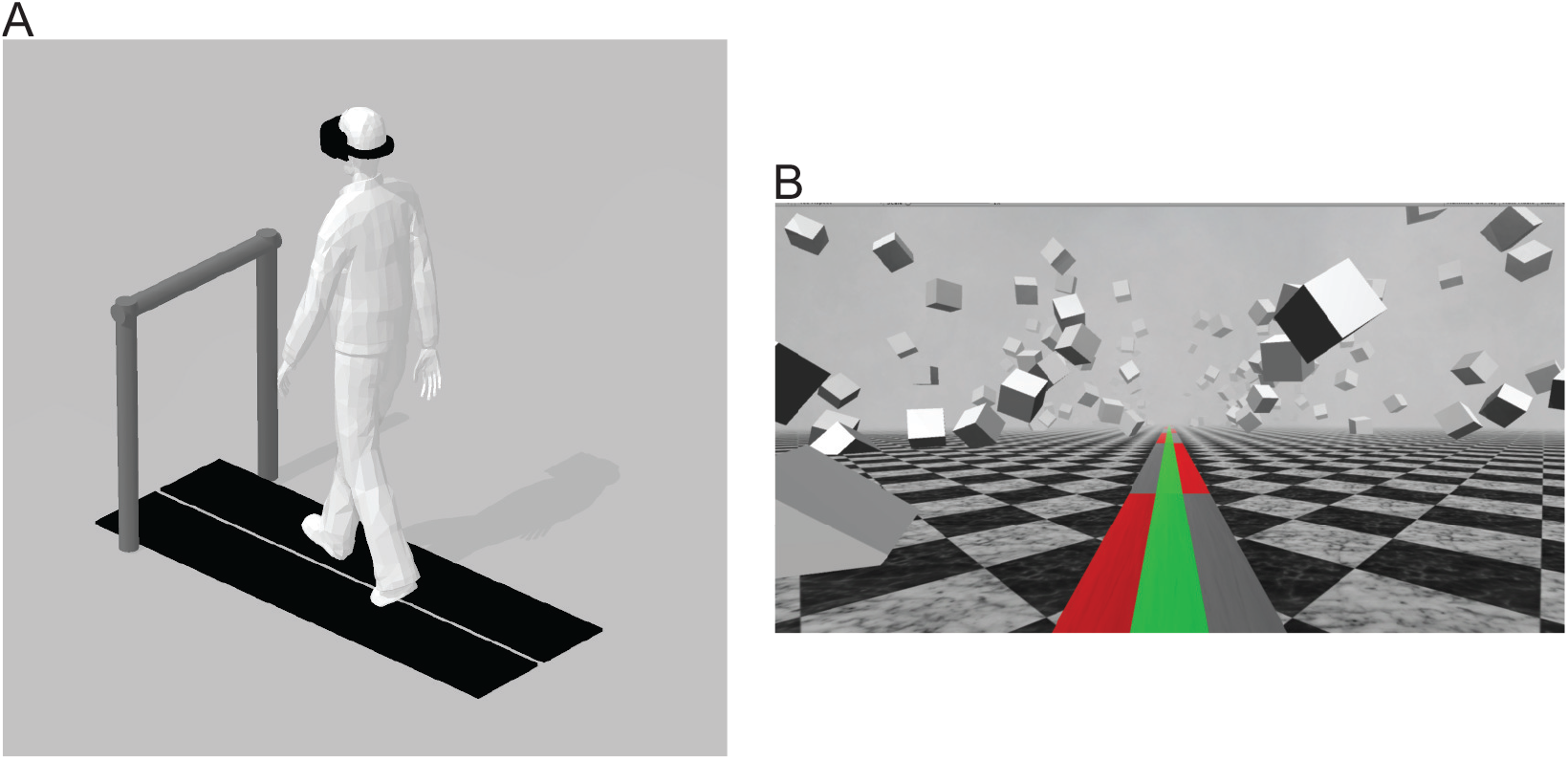
(A) Subjects walked on a self-paced treadmill, wearing a head-mounted display, while harnessed to a body-weight support system (not shown). The front handrail was used to assist in starting and stopping the selfpaced treadmill comfortably. (B) The virtual reality scene displayed in the Oculus Rift. Subjects were instructed to walk on the green pathway and avoid stepping in the red *No-Step* zones. If the subject needed to, they were able to step in the gray *Neutral* zones.

The virtual environment was designed and implemented in Unity3d (Unity Technologies, San Francisco, CA, USA) and consisted of a tiled marble floor with floating cubes randomly distributed in a volume 0-10 m above the floor, 2-17 m to each side from the midline, and infinitely into the distance providing peripheral visual depth and texture. On top of the marble floor was a pathway with three colored lanes of equal width. These three virtual lanes were aligned with the treadmill belts and, in combination, fully covered the width of the treadmill (1.10 m, see Figure 1B). The middle lane was colored green. The left and right lanes alternated between gray and red, infinitely into the distance, each block with a length of 10 m. Instructions to the subjects were “Walk on the green path. Please do not step onto the red zones. It is ok to step onto the gray zones if you have to.” We will refer to the red areas as “No-Step” and the gray areas as “Neutral” zones.

After explaining the experiment, obtaining consent and placing markers and EMG sensors, subjects first walked for 15 minutes to adapt to the self-paced treadmill with the Oculus Rift. The instructions for starting the treadmill were to hold the handrail until the treadmill reached a comfortable pace, then let go and walk normally. During the adaptation we told the subjects that we would now perturb their sense of balance by use of the galvanic vestibular stimulation (GVS), and asked them to cope with this perturbation “normally” and keep walking forward. Data collection blocks consisted of alternating phases of metronome and stimulus. During metronome phases, lasting 30 s, we provided an auditory metronome at 100 bpm and asked subjects to use this as an “approximate guideline” for their footsteps, both during metronome and stimulus phases. During stimulus phases, lasting 120 s, we turned the metronome off, and subjects received intermittent balance perturbations (details below). Data were collected during stimulus phases. Each subject performed four blocks of walking, each block consisting of five metronome and five perturbed phases, always starting with metronome phases, for a total of 12.5 min per block. After each block, the treadmill was turned off and subjects were offered a break. This protocol was implemented in a custom Labview program that sent the head position and treadmill speed to the Unity computer via UDP and saved the GVS current and treadmill speed at 100 Hz.

Forty-four reflective markers were placed on the subject, using the Plug-in Gait marker set (Davis III et al., 1991), with six additional markers on the anterior thigh, anterior tibia, and 5th metatarsal of each foot. Marker positions were recorded at 200 Hz using a Qualisys motion capture system with 13 cameras. Ground reaction forces and moments were collected at 1000 Hz from both sides of the split-belt treadmill and transformed into a common coordinate frame to calculate whole-body center of pressure (CoP).

The vestibular perturbations were triggered on heel-strikes of either foot. The direction of the fall, determined by the polarity of the current, was always chosen in the direction of the foot triggering the stimulus. We identified heel-strikes as the local maxima of forward progression of the heel marker. Due to the alternation of the zones in the outer lanes, this could provide a sensation of falling either towards a *No-Step* or towards a *Neutral* zone. We also included a sham condition with no stimulus as an unperturbed reference, and randomly chose each trigger to be GVS or sham, with equal likelihood. GVS Perturbations consisted of a 1 mA current passed between two round electrodes (3.2 cm diameter, Axelgaard Manufacturing 103 Co., Ltd, Fallbrook, CA, USA) placed on the mastoid processes for 1 s, inducing a feeling of falling to the side. The perturbation could only be triggered in the middle third between a No Step and a *Neutral* zone to avoid possible confusion near the zone switch lines, and was randomized to trigger 1-4 steps after entering the valid inner area. To allow the effects of the perturbation to wash out, further triggers were blocked for at least 10 steps.

### 2.2. Data analysis

We low-pass filtered force plate data with a 4th order Butterworth filter at a cut-off frequency of 20 Hz and kinematic data with a 4th order Butterworth filter with a cutoff frequency of 10 Hz. We calculated joint angle data from the marker data, based on a geometric model with 15 segments (pelvis, torso, head, thighs, lower legs, feet, upper arms, forearms, hands) and 38 degrees of freedom (DoF) in OpenSim (Delp et al., 2007; Seth et al., 2018) using an existing base model (Zajac et al., 1990). We calculated the body center of mass (CoM) trajectories based on this model. To normalize the EMG trajectories, we divided each signal by the average over all control strides for each channel and subject.

We normalized the data between heel-strikes to 100 time steps. To estimate the motor response to the perturbations, we subtracted the average of the unperturbed sham triggers within each subject for all data. Thus all of results will be presented as a difference from control, referred to as a response.

To improve the estimate of the foot placement response to the stimulus, we fitted a linear regression model relating the foot placement changes for each subject to the changes of lateral position and velocity of the CoM at midstance using the control data (Wang and Srinivasan, 2014). Then for each stimulus step, we used this model to estimate the expected foot placement change based on the CoM state, and subtracted this from the observed foot placement change, resulting in an estimate of the foot placement change due to the vestibular stimulus (Reimann et al., 2017). We will refer to this model-based estimate as model-corrected foot placement change.

The experimental design contains two distinct stimulus conditions, where each stimulus could induce a fall sensation either towards a *No-Step* zone or towards a *Neutral* zone. We will refer to these two conditions simply as *No-Step* or *Neutral* stimuli from here on. Spatially, the fall was always towards the triggering leg, which could be either the left or the right. After processing and filling gaps in the kinematic data, we were left with 1211 *No-Step* and 1207 *Neutral* stimuli.

Subjects were able to successfully maintain balance while walking on the self-paced treadmill, with no instances of stepping off the treadmill or making use of the body-weight support system. Occasionally subjects violated the task by stepping into the *No-Step* zone. Across all subjects, 64 such violations occurred during or after *No-Step* stimuli and 47 during or after *Neutral* stimuli. We removed these stretches of data from further analysis, which left 1147 *No-Step* and 1160 *Neutral* stimuli.

### 2.3. Outcome Variables

We expected that the effect of the vestibular perturbations would be different for fall stimuli toward *No-Step* zones compared to those toward *Neutral* zones. Our first hypothesis was that the whole-body CoM excursion would be larger for *No-Step* vs. *Neutral*. We further hypothesized that responses in the foot placement mechanism would be smaller, and responses in the ankle roll and push-off mechanism would be larger for *No-Step* vs. *Neutral*. Here we specify main and secondary outcome variables we calculated to test these hypotheses statistically.

For the overall balance response, we used (i) the Δ shift of the whole-body CoM at the end of the fourth step and (ii) the Δ velocity at the end of the second post-stimulus step, based on previous results showing that the maximal changes are near these times (Reimann et al., 2018).

For the ankle roll mechanism, we analyzed four variables related to the stance leg lateral ankle activation: (iii) the Δ CoP-CoM distance integrated over the first post-stimulus single stance. (iv) the Δ stance leg ankle eversion/inversion angle integrated over the first post-stimulus single stance. (v) the Δ stance leg peroneus longus EMG integrated over the first post-stimulus single stance. (vi) the Δ stance leg tibialis anterior EMG integrated over the first post-stimulus single stance.

For the foot placement mechanism, we analyzed five variables related to the first post-stimulus swing leg heel strike: (vii) the foot placement, defined as the Δ swing leg heel position relative to the trigger leg heel position at swing leg heel strike. (viii) the model-corrected foot placement, defined as the difference between the measured foot placement value and the foot placement value predicted based on the position and velocity of the CoM at mid-stance using the linear model (see above). (ix) the Δ trigger leg knee internal/external rotation angle at swing leg heel-strike. (x) the Δ swing leg hip internal/external rotation angle at swing leg heel-strike. (xi) the Δ swing leg hip abduction/adduction angle at swing leg heel-strike.

For the push-off mechanism, we analyzed two variables related to the push-off of the trailing leg before and during the second post-stimulus double stance: (xii) the Δ plantarflexion angle integrated over the second post-stimulus double stance phase. (xiii) the Δ medial gastrocnemius EMG of the stance leg integrated over the first post-stimulus swing phase.

### 2.4. Statistical analysis

We confirmed the assumptions of normality and homoscedasticity by visual inspection of the residual plots for the variables related to foot placement, lateral ankle, and pushoff mechanisms. Our primary focus of analysis is the whole-body CoM shift and the kinematic, kinetic, and electromyographical variables associated with the ankle roll, foot placement and push-off balance mechanisms.

To test our hypotheses about whether humans use the balance mechanisms differently across stimulus directions, we used R (R Core Team, 2013) and lme4 (Bates et al., 2009) to perform a linear mixed effects analysis. For each outcome variable, we fitted a linear mixed model and performed an ANOVA to analyze the use of the mechanisms and interaction of stimulus direction, using Satterthwaite’s method (Fai and Cornelius, 1996) implemented in the R-package lmerTest (Kuznetsova, 2017). As fixed effects, we used the direction of the perturbation towards a *No-Step* or a *Neutral* zone. As random effects, we used individual intercepts for subjects. Table 1 displays the results for this statistical test on the outcome variables.

**Table 1:** Results of the linear mixed model ANOVA. Bold face text indicates statistically significant differences between perturbations toward *No-Step* and *Neutral* zones.

| Variable | Num. Df | Den. Df | F | p |
| --- | --- | --- | --- | --- |
| $\Delta$ CoM | 1 | 1922.3 | 145.38 | <**0.0001** |
| $\Delta$ CoM Velocity | 1 | 1921 | 31.105 | <**0.0001** |
| $\int \Delta$ CoP-CoM | 1 | 1928.7 | 0.0946 | 0.7585 |
| $\int \Delta$ Ankle Eversion | 1 | 1930.1 | 2.4456 | 0.118 |
| $\int \Delta$ Peroneuous Longus EMG | 1 | 1924.8 | 2.657 | 0.1033 |
| $\int \Delta$ Tibialis Anterior EMG | 1 | 1928.7 | 3.8821 | **0.04895** |
| $\Delta$ Foot placement | 1 | 1925.4 | 0.9193 | 0.3378 |
| Model-Correct Foot placement | 1 | 1921.4 | 29.15 | <**0.0001** |
| $\Delta$ Knee Rotation | 1 | 1924 | 0.0637 | 0.8008 |
| $\Delta$ Hip Rotation | 1 | 1922.3 | 3.593 | 0.05817 |
| $\Delta$ Hip Adduction | 1 | 1923.6 | 0.1343 | 0.714 |
| $\int \Delta$ Ankle Plantarflexion | 1 | 1927 | 2.789 | 0.09508 |
| $\int \Delta$ Medial Gastroc EMG | 1 | 1924.7 | 4.6848 | **0.03055** |

## 3. Results

Table 1 shows the results of the linear mixed model ANOVA. Statistically significant differences between *No-Step* and *Neutral* are marked in bold.

We observed a whole-body balance response in both the *No-Step* and *Neutral* conditions. Figure 2A shows the effect of the balance response in the center of mass (CoM) shift in both conditions. The average CoM trajectories for the two conditions start to separate around the third post-stimulus step. As hypothesized, the effect of the perturbation is stronger in the *No-Step*. This difference is reflected in the CoM velocity results, shown in Figure 2B. Both of these differences are statistically significant (see Table 1).

**Figure 2:**
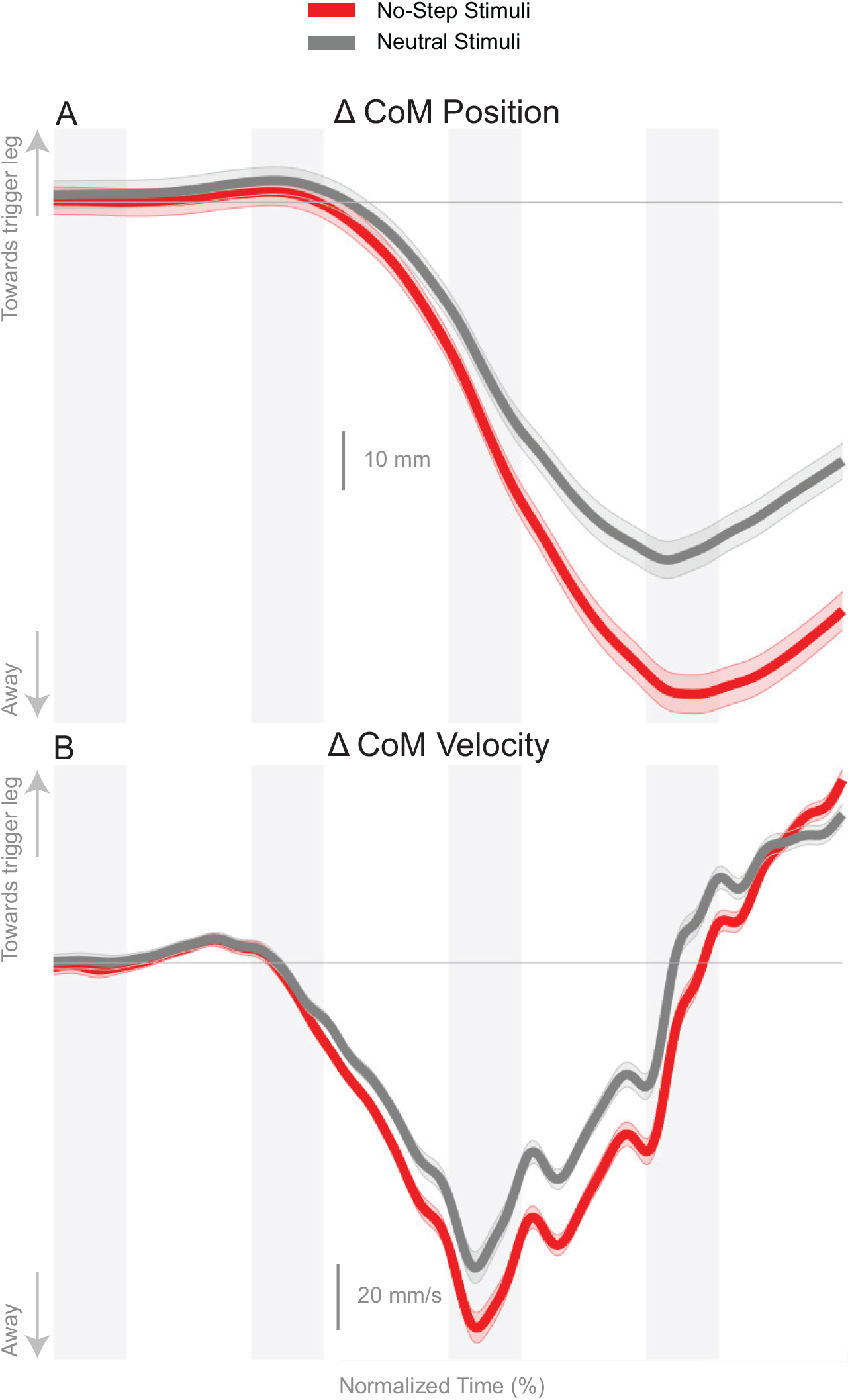
Changes in response to the balance perturbation in the medial-lateral CoM position (A) and velocity (B). Curves start at the heel-strike triggering the stimulus and span four steps, ending at heel-strike of the same foot. Red curves correspond to fall stimuli toward the *No-Step* and gray curves to fall stimuli toward the *Neutral* zone. The gray vertical bars correspond to double stance phases and the white areas in between to single stance phases. Data shown are changes relative to the unperturbed steps, with shaded areas around the means giving the 95% confidence intervals.

Figure 3 shows the displacement of the CoP relative to the CoM. The CoP shifts in the direction of the perceived fall in both conditions, as expected. There appears to be no difference in the response to the perceived fall between the two conditions during the single stance period, though after the first post-stimulus heel-strike the average trajectories begin to diverge.

**Figure 3:**
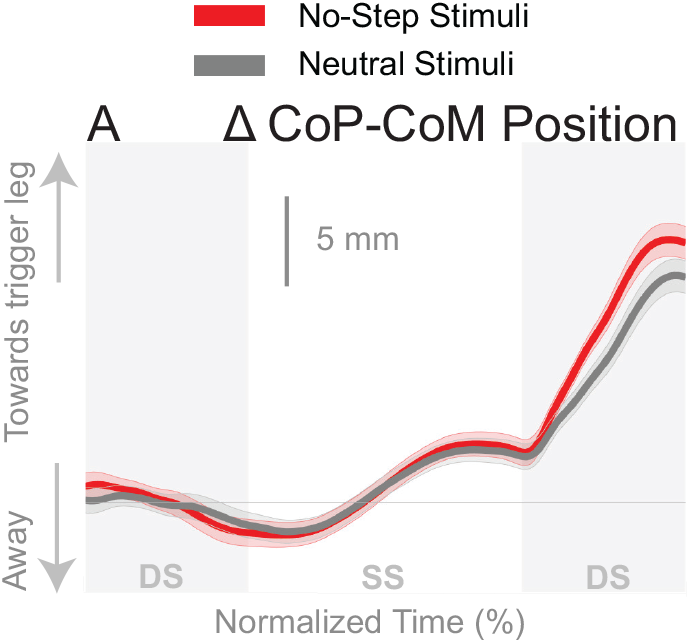
Changes in the medial-lateral CoP-CoM displacement in response to the perturbations. Curves start at the heel strike triggering the stimulus and end at the triggering foot toe-off. Red curves correspond to fall stimuli toward the *No-Step* and gray curves to fall stimuli toward the *Neutral* zone. The vertical shaded areas correspond to double stance, the white area between to single stance. Data shown are changes relative to the unperturbed steps, with shaded areas around the means giving the 95% confidence intervals.

### Ankle Roll

The shift of the CoP and CoM are generated by a temporally coordinated set of balance mechanisms. Due to the stimulus occurring on heel strike, the first available response is in the ankle roll mechanism. Figure 4 shows the kinematic and electromyographic response of the lateral ankle mechanism. Figure 4A displays a triggering leg inversion during the first post-stimulus step relative to the unperturbed pattern. The ankle inversion change trends higher towards the end of single stance, but is not statistically different. Figure 4B displays the peroneous longus muscle activity, an ankle everter, and Figure 4C displays the tibialis anterior, an ankle inverter. Together, the changes in activation of these two muscles provide evidence the lateral ankle mechanism is actively generated. A decrease in peroneous longus activity and an increase in tibialis anterior activity yield an ankle inversion.

**Figure 4:**
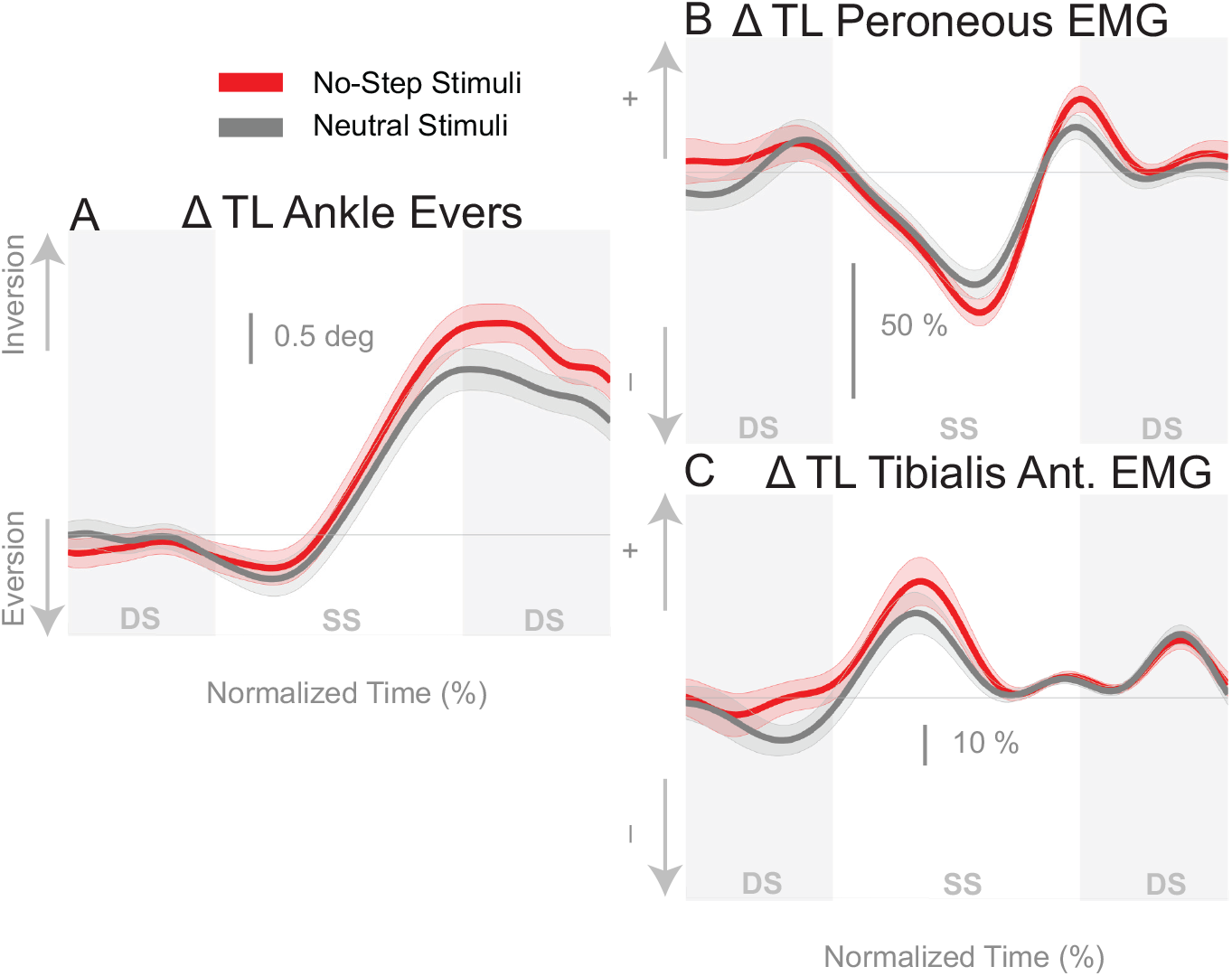
Variables illustrating the use of the lateral ankle mechanism. (A) The joint angle that contributes to the CoP-COM changes; and (B,C) the EMG that contributes to joint angle changes. Curves start at the heel strike triggering the stimulus and end at the triggering foot toe-off. Red curves correspond to fall stimuli toward the *No-Step* and gray curves to fall stimuli toward the *Neutral* zone. The vertical shaded areas correspond to double stance, the white area between to single stance. Data shown are changes relative to the unperturbed steps, with shaded areas around the means giving the 95% confidence intervals.

### Foot Placement

Figure 5A shows that the foot placement was used in both conditions, as evidenced by the shift of the average response in the direction of the perceived fall. The difference in foot placement response was not statistically significant between the *No-Step* and *Neutral* conditions (1). The better estimate of the model-corrected foot placement change, however, shows a stronger difference between zone conditions that did reach statistical significance (Table 1), as seen in Figure 5B. Contrary to our expectation, the foot placement response to stimuli towards the *No-Step* zone was larger than the response to stimuli towards the *Neutral* zone, rather than smaller. Figure 6 shows the kinematic variables related to the foot placement. The the swing leg hip adducts, and internally rotates, and the trigger leg knee internally rotates, for both conditions. The combination of these three joint angle changes yield the foot placement change in the direction of the fall stimulus, displayed in Figure 6A. None of these kinematic measures are significantly different between conditions (Table 1).

**Figure 5:**
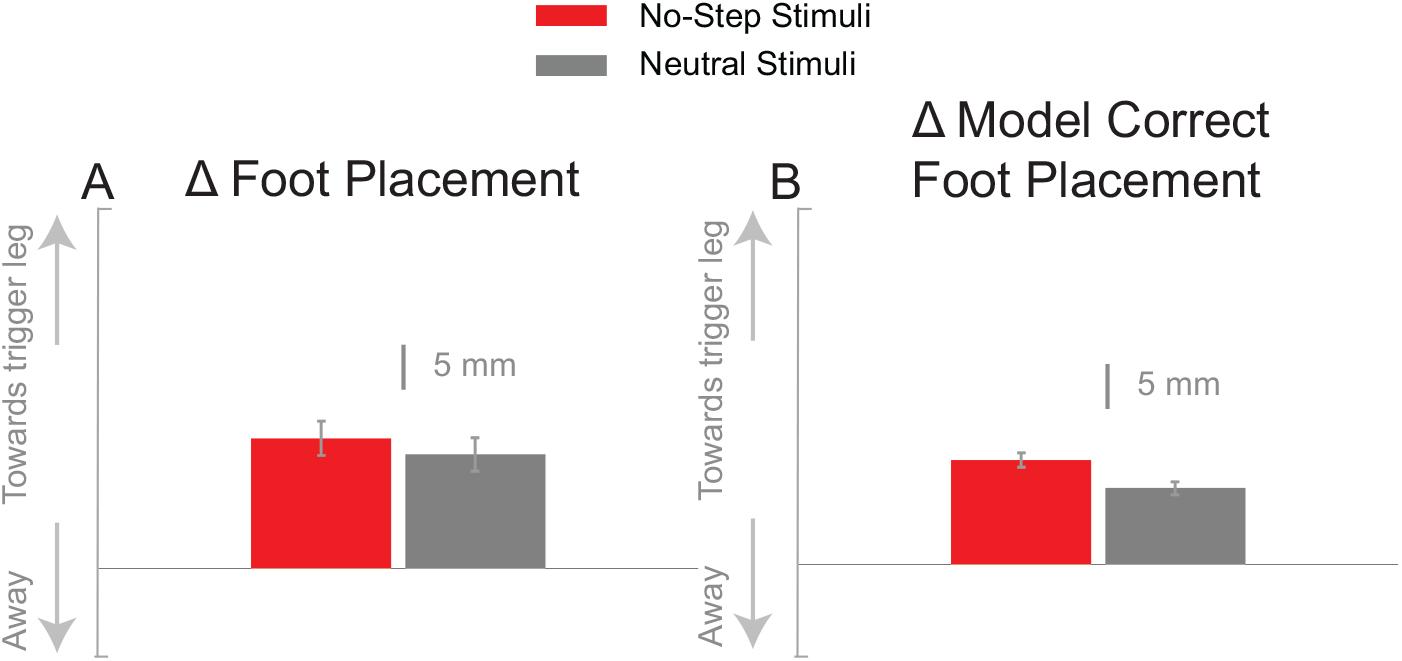
The foot placement response (A) and the model-corrected foot placement (B) at the first post-stimulus step. Bars indicate the mean and the errors the 95% confidence interval.

**Figure 6:**
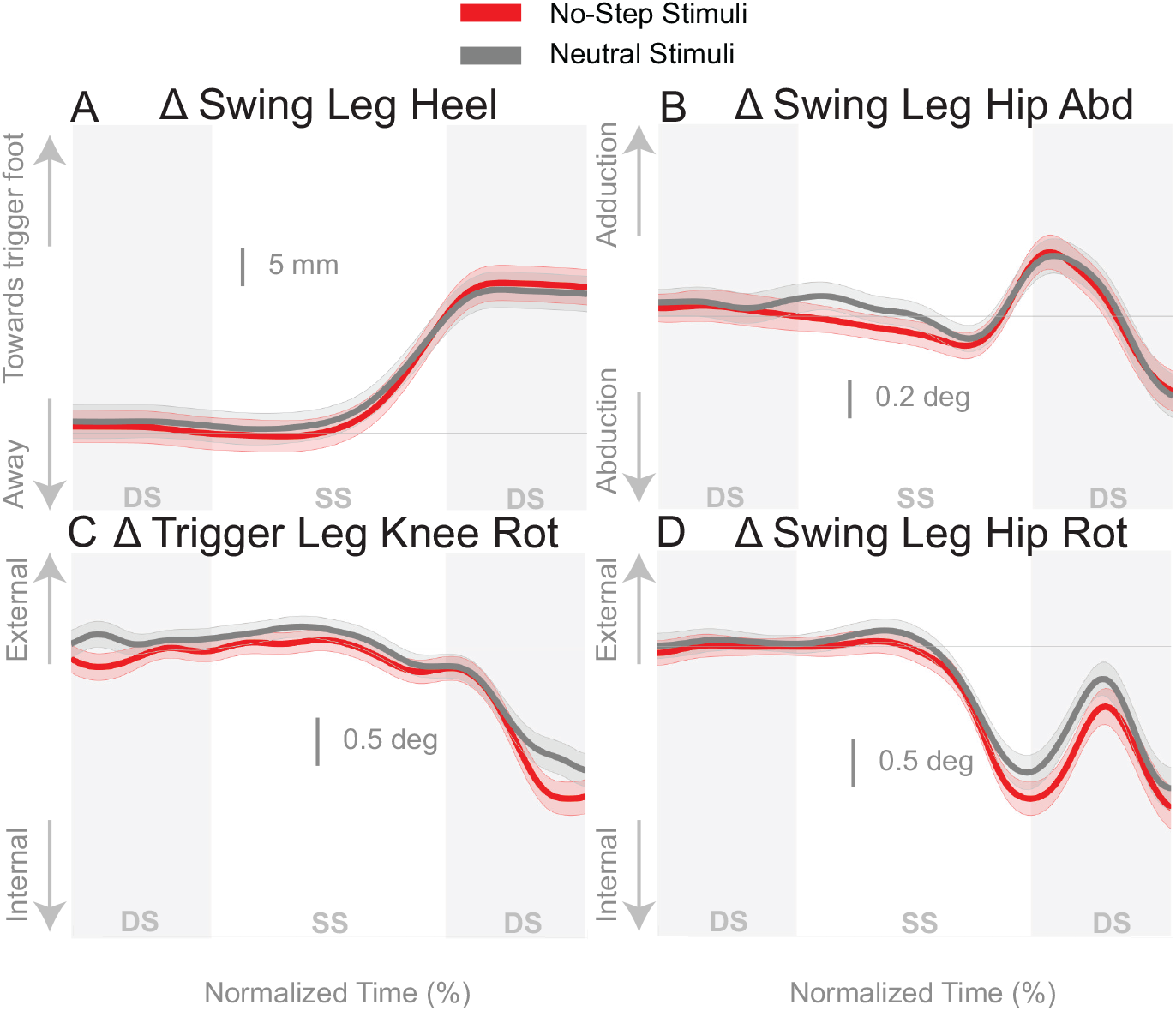
Variables illustrating the use of the change of the foot placement mechanism over time. Changes in response to the balance perturbation in the heel position (A), and joint angles (B,C,D) that contribute to change in heel position. Curves start at the heel strike triggering the stimulus and end at the triggering foot toe-off. Red curves correspond to fall stimuli toward the *No-Step* and gray curves to fall stimuli toward the *Neutral* zone. The vertical shaded areas correspond to double stance, the white area between to single stance. Data shown are changes relative to the unperturbed steps, with shaded areas around the means giving the 95% confidence intervals.

### Push-off

Figure 7 shows that the push-off mechanism was also used to respond to the balance perturbation, with an increased plantarflexion and increased medial gastrocnemius activity for both conditions. According to Table 1, the integrated plantarflexion angle change quantifying the kinematic resposne did not significantly differ between conditions, but the medial gastrocnemius activity did differ significantly, with more medial gastrocnemius activity in the *No-Step* condition.

**Figure 7:**
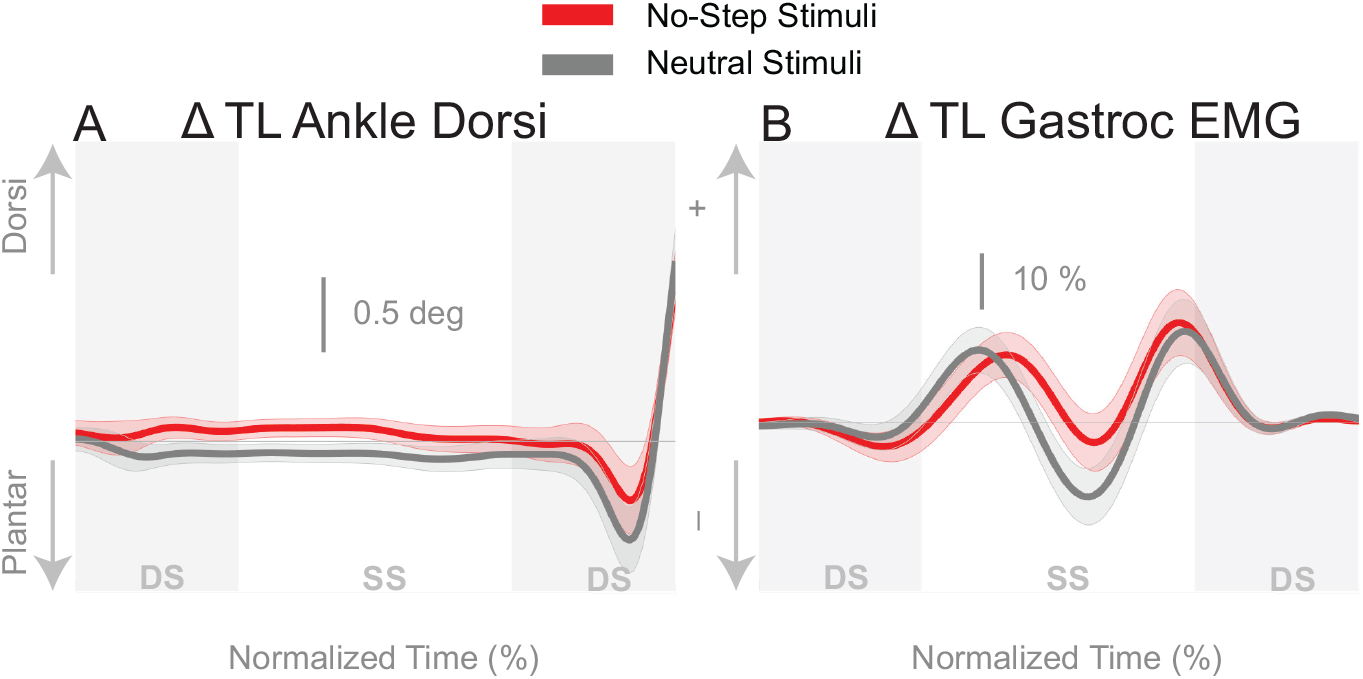
Variables illustrating the use of the push-off mechanism. Changes in response to the balance perturbation in ankle dorsiflexion (A), and the medial gastrocnemius EMG (B). Curves start at the heel strike triggering the stimulus and end at the triggering foot toe-off. Red curves correspond to fall stimuli toward the *No-Step* and gray curves to fall stimuli toward the *Neutral* zone. The vertical shaded areas correspond to double stance, the white area between to single stance. Data shown are changes relative to the unperturbed steps, with shaded areas around the means giving the 95% confidence intervals.

## 4. Discussion

This experiment served as the first attempt in identifying how environmental constraints modify the use of the different balance mechanisms in walking. We expected virtual constraints would alter the use of the balance mechanisms. The virtual environment, shown in Figure 1B, paired with instructions not to step onto the red *No-Step* zones, was used to determine if environmental constraints alter the use of the balance mechanisms. We found that this paradigm produced subtle changes in the balance mechanisms that led to a robust difference in the whole-body CoM displacement.

We observed a separation of the CoM movement in the third and fourth steps following the stimulus onset. There was more of an overall shift of the CoM when the balance perturbation was towards the *No-Step* zone compared to the *Neutral* zone. We expect the CoM shift to be a result of the use of the balance mechanisms. Although there were no strong differences between the zone conditions for any of the balance mechanisms, we observe subtle trends in each mechanism that are consistent between multiple variables associated with each balance mechanism.

The lateral ankle mechanism appears to be consistently larger in the *No-Step* condition, with larger ankle inversion, decreased peroneous muscle activity, and increased tibialis anterior activity, shown in Figure 4. Only the tibialis anterior muscle activity showed a significant difference between the *No-Step* and *Neutral* conditions, but in combination the evidence suggests that the ankle roll mechanism was activated to a greater extent when a perturbation induced a fall toward the *No-Step* zone. The foot placement mechanism also shows trends to support consistently larger use in the towards *No-Step* Zone condition, contrary to our hypothesis. Only the model corrected foot placement showed a significant difference between perturbation directions for evidence of a greater use of the foot placement mechanism. Regardless, this difference, in combination with the subtle trends of kinematics shown in figures 6 and Figure 5A, provide evidence to suggest increased use of the foot placement mechanism when the perturbation induces a fall toward the *No-Step* zone. The push-off provides less compelling evidence to suggest a difference between conditions, despite a difference in medial gastrocnemius activity. Figure 7A shows a greater plantarflexion occurs in *Neutral* zone perturbations, but figure 7B shows the muscle activity is trending lower relative to *No-Step* zone perturbations. The combination of the subtle differences in the lateral ankle and the foot placement mechanisms in the first step post-stimulus likely generate the CoM shift we observe in the third and fourth steps post-stimulus. This is evidence that humans can indeed alter the mechanisms they use to maintain balance, particularly in the medial-lateral direction, based on perceived environmental constraints.

The modification of the balance response in the presence of an environmental constraint indicates substantial influence from central, supraspinal components. The balance response was altered, despite randomization of perturbation direction, suggesting a descending command that influences the execution of the balance response. However, unlike previous research regarding supraspinal central sets in balance control (Horak et al., 1989), here we observe a direction dependent adjustment instead of amplitude or velocity. If a direction dependent central set were in place, a primed activation of the balance mechanisms can push the body further from the undesired location (*No-Step* zone). In general, the external constraint imposed leads to a cognitive bias that effects the response to the perturbation.

Despite the changes we see in the balance mechanisms dependent on the stimulus direction, the observed foot placement response was contrary to our hypothesis. We expected a decreased foot placement response during *No-Step* stimuli, but instead observed an increased use of the foot placement mechanism. Multiple factors could contribute to this result: 1) the participants did not have great perception of their lower limbs while totally immersed in virtual reality, 2) the green pathway was wide enough to make a foot placement while still abiding by the rules of the task, or 3) the participants did not perceive the task as having a high degree of risk (i.e. no consequences for stepping in area). We discuss these factors and possible solutions in the following paragraphs.

We intentionally did not provide any visual or auditory feedback of whether the individual was completely adhering to the protocol (i.e. never stepping in the red pathway). We opted for a minimalist development with no avatar or audio feedback. It is arguable that the presence of a virtual avatar would help the perception of the participant’s own body in space. Previous research has shown motor adaptation in upper limb experiments in the presence of an avatar (Bourdin et al., 2019). However, studies about gait kinematics are still needed to understand the specific components of virtual reality immersion and their possible benefits (Ferreira dos Santos et al., 2015).

Future experiments should look to improve the perception or effect of the environmental constraint. In the example of walking along a cliff, the cost of not abiding by the constraints is high; a fall off the cliff. This urgency does not exists in the current paradigm. We avoided the confounding factor of fear by creating a mundane environment and provided instructions to simply avoid stepping on a particular color. Fear and anxiety are known to modify behavior (Raffegeau et al., 2020) and spinal reflexes when exposed to increased heights (Sibley et al., 2007). Fear may be one method to provide an incentive to abide by such a constraint. Another strategy to improve the efficacy of the paradigm could be to narrow the path of normal walking or decrease the distance between the no-step zone and walking path. However, we would likely not want to constrain the path to the extent of narrow beam walking (Peterson and Ferris, 2018), as this form of walking entirely eliminates the ability to use the foot placement mechanism for the control of balance in the medial-lateral direction (Otten, 1999). It may also be possible to improve engagement through gamification of the task. Positive feedback for avoiding the no-step zone or negative feedback for stepping in the no-step zone may improve the engagement in this paradigm. Although virtual reality is still in its infancy, many researchers have attempted to modify behavior through virtual reality in neurological populations (Felsberg et al., 2019). Further, a handful of researchers are attempting to modify and improve walking performance through gamification (Eggenberger et al., 2016; Schättin et al., 2016; Adcock et al., 2020).

This knowledge and paradigm will be particularly helpful for individuals that suffer from an injury or condition that prevent the use of a specific balance mechanism. For example, people with diabetic neuropathy or Parkinson’s disease might refrain from using the ankle roll mechanism due to diminished proprioception. A paradigm similar to the one described here could be used to enhance the use of the ankle roll mechanism. We used vestibular stimulation to elicit balance responses, but similar elicitations could be provoked by mechanical perturbations (Banala et al., 2010; Acasio et al., 2017).

## 5. Conclusion

We created a task for walking in a virtual environment that can be used to study the effects of environmental constraints on the control of balance during locomotion. The results indicate that the neural control of balance changes in the presence of environmental constraints. More research is needed to refine the paradigm to determine how exactly the balance mechanisms change in the presence of environmental constraints. If we can predict which balance mechanisms people will use in specific situations, we can use this to intervene in the automatic balance response through alteration of the environment. Such a paradigm may be an avenue to target and rehabilitate specific balance mechanisms and strategies.

## Author contribution statement

TF, FDS, HR, JJ designed the experiment. TF, HR, and SD performed the experiment, TF, HR, SD analyzed the data. TF, FDS, HR, SD, and JJ wrote the manuscript.

## Notes

#### Summary of Updates

There was a typo in the discussion where I intended to write a brief paragraph. I have now removed the typo and added a paragraph.

